# Reprogrammed *E. coli* for secretion of thermophilic cellulase cocktail for seawater-compatible lignocellulosic bioprocessing

**DOI:** 10.1101/2025.07.27.664954

**Authors:** Mani Gupta, Aniket Sarkar, Suksham, Debdeep Chatterjee, Supratim Datta

## Abstract

The development of efficient cellulase systems is crucial for sustainable lignocellulosic biorefining. In this study, we engineered Escherichia coli to co-express and secrete a thermophilic cellulase cocktail consisting of a Bacillus licheniformis processive endoglucanase (H1AD14), a Chaetomium thermophilum cellobiohydrolase (G0SAE6), and a glucose-tolerant engineered β-glucosidase B8CYA8tm (V169C/E173L/I246A B8CYA8) from Halothermothrix orenii. Fusion with the AnsB secretion tag enabled extracellular yields of 0.2-0.4 g/L for each enzyme, bypassing costly purification while retaining activity. The secretion was confirmed via immunoblotting using anti-His antibody, and catalytic activity was validated through enzymatic assays. The absence of E. coli native periplasmic maltose-binding protein (detected with anti-MBP antibody) in the extracellular medium ruled out cell lysis, confirming protein secretion. Using the Golden Gate assembly, we constructed and systematically evaluated synergistic enzyme combinations for hydrolytic performance. By optimizing secretion conditions, including media, inducer concentration, and incubation temperature, the total yield of secreted enzyme cocktail increased from 0.31 g/L to 0.74 g/L. The secreted cocktail effectively hydrolysed diverse lignocellulosic substrates and exhibited 100 % activity in seawater, demonstrating potential for freshwater-independent biorefining. The work addresses key economic and environmental challenges in biomass conversion by integrating secretion engineering, thermostability, and seawater tolerance into a single scalable system.

## Introduction

The hydrolysis of lignocellulosic biomass by cellulases presents a sustainable route for biofuel production, offering an alternative to fossil fuels^1^. Efficient biomass degradation requires synergistic action of three key cellulases-Endoglucanase (EG), cellobiohydrolase (CBH), and β-glucosidase (BG)^2-4^. However, natural enzyme systems often lack the thermostability and inhibition tolerance needed for industrial applications. *Trichoderma reesei*, a well-known cellulase producer, suffers from insufficient β-glucosidase activity^5, 6^, necessitating supplementation from other microbial sources. Since no single organism naturally produces an optimal enzyme cocktail, the selection and engineering of high-performance cellulases are critical for efficient biomass conversion. Thermotolerant cellulases offer distinct advantages, as they enhance hydrolysis efficiency by operating at high temperatures, resisting inhibitors, and reducing microbial contamination^7, 8^. High-temperature processing also improves substrate solubility and increases enzyme accessibility to cellulose fibers. These properties make thermophilic enzymes particularly attractive for cost-effective biorefining^9, 10^.

Extracellular cellulase secretion from the host-microbe has several advantages over intracellular protein expression. A significant benefit of Escherichia coli is the reduction in enzyme degradation due to the low levels of *E. coli* proteases in the extracellular medium compared to those in the cytoplasm. The lower extracellular protein concentrations minimize aggregation and inclusion body formation. Secretion also simplifies downstream processing by enabling direct use of culture supernatants, bypassing costly purification steps. While *E. coli* typically retains recombinant proteins in the cytoplasm or periplasm, engineering of secretion pathways using signal peptides or secretory proteins PelB, OmpA, OsmY, YebF, and TorA^11-15^. Recent studies suggest that a typical secretion pathway can significantly improve extracellular protein production, enabling direct use of the culture medium for industrial applications eliminating costly purification steps.

*E. coli*’s rapid growth and genetic tractability make it an ideal host for recombinant cellulase production. Although lacking post-translational modifications found in eukaryotic systems, it can facilitate disulfide bond formation and multi-enzyme co-expression, which improves folding, reduces inclusion bodies, and boosts catalytic efficiency^16-19,20^. For instance, co-expressing xylanase and laccase in *Pichia pastoris* increased total degradation activity by 44 %^21^. Kumar et al. engineered a multi-enzyme construct containing laccase, pectate lyase, and endoxylanase in *E. coli*, significantly enhancing the saccharification of orange peel compared to xylanase alone^22^. In another study, Zhang et al. co-expressed xylanase, β-xylosidase, and feruloyl esterase for efficient hemicellulose degradation^20^. These findings underscore the potential of co-expressing enzyme cocktails for biomass processing.

To maximize applicability, we developed a thermostable cellulase cocktail comprising a glucose-tolerant BG (B8CYA8tm, V169C/E173L/I246A mutant) with high activity and stability, a cellobiohydrolase (G0SAE6) from *Chaetomium thermophilum*, and a processive endoglucanase (H1AD14, GH9) from *Bacillus licheniformis* with broad substrate specificity^25-23^. We incorporated the *E. coli* L-asparaginase II (AnsB) signal peptide, an underexplored secretion tag for high-yield recombinant protein export. Compared to the widely used fusion protein YebF^14^, and signal peptides like OmpA^24-26^, phoA^27^, and pelB^28^, the AnsB tag enabled significantly higher secretion yields. Another major bottleneck in industrial-scale lignocellulosic biofuel production is the high freshwater demand, which limits sustainability and scalability^29-31^. To address this, we demonstrated that this cellulase cocktail retains full hydrolytic activity in unprocessed seawater, eliminating the freshwater in biomass saccharification. This breakthrough, combined with the modular design of our secretion platform, provides a scalable solution for cost-effective, environmentally compatible biofuel production.

## Materials and methods

### Chemicals

All chemicals used in this study were of reagent grade. DNA polymerase (Q5 High-Fidelity DNA polymerase), restriction endonucleases, Golden Gate Assembly Master Mix, HiFi DNA Assembly Master Mix, phosphatase, and DNA ligase were from NEB (MA, USA). dNTP Mix and DNA marker were from Thermo Scientific (Mumbai, India). Primers were synthesized by Barcode Biosciences (Bangalore, India). The commercial cellulase, chemicals, and substrates like carboxy methyl cellulose (CMC) and Avicel were purchased from Sigma-Aldrich (St Louis, MO, USA). The chromogenic substrates, like *p*-nitrophenyl β-d-glucopyranoside (*p*NPGlc), were purchased from TCI (Tokyo, Japan). FemtoLUCENT™ PLUS-HRP kit was purchased from G Biosciences (Saint Louis, Missouri, USA). Plasmid miniprep and agarose gel purification kits were bought from Qiagen (Hilden, Germany A centrifugal concentrator with a 30 kDa molecular weight cut-off membrane was purchased from Merck Millipore (Bangalore, India).

### Bacterial strains, plasmids, and growth conditions

The bacterial strains and plasmids used in this study are listed in SI Table 2. *E. coli* DH5α was used as the host strain for standard cloning procedures, while *E. coli* BL21 (DE3) served as the expression host for recombinant protein production using EcoFlex plasmids (pTU1A, pTU1B, pTU1C, pTU2A, and pTU2B). ^32^ Strains were cultured aerobically in Luria-Bertani (LB) medium at 30 °C or 37 °C, with ampicillin (100 μg/mL) or chloramphenicol (25 μg/mL) added when necessary for plasmid selection and maintenance. For recombinant strain cultivation and target protein production various media formulations were used, including LB medium (peptone 10 g/L, yeast extract 5 g/L, NaCl 10 g/L), SB medium (peptone 32.0 g/L, yeast extract 20.0 g/L, NaCl 5.0 g/L), 2× YT medium (peptone 16.0 g/L, yeast extract 10.0 g/L, NaCl 5.0 g/L), and SOB medium (peptone 20.0 g/L, yeast extract 5 g/L, NaCl 0.5 g/L, KCl 0.2 g/L, MgCl₂ 1.0 g/L, MgSO₄ 1.2 g/L). These optimized media formulations were selected based on their ability to support high cell density growth and maximize recombinant protein yield.

### Construction of enzyme expression plasmids

The open reading frames (ORFs) of EG, CBH, and BG were PCR amplified using primers EG_FP/EG_RP, CBH_FP/CBH_RP, and BG_FP/BG_RP, respectively (all primers used in cloning are listed in SI Table 1). These ORFs were then cloned into the pBP-ORF plasmid using *Bsa*I restriction sites. To incorporate the AnsB secretion, the AnsB sequence was PCR amplified with AnsB_FP/AnsB_RP primers and inserted into the pBP-lacZ vector from the EcoFlex kit using *Nde*I and *Sph*I restriction sites. Next, the ORFs were assembled into Level 1 expression plasmids (pTU-1A, pTU-1B, and pTU-1C) using the Golden Gate assembly method with the EcoFlex Kit^32^. The assembly combined pBP-T7 promoter, pBP-PET RBS, pBP-AnsB_tag, the enzyme ORFs cloned in pBP-ORF, and pBP-BBa0015 terminator. Successful Level 1 assembly was validated through colony PCR and sequencing. Level 2 assembly was performed to make these four combinations (i) AnsB_EG + AnsB_CBH, (ii) AnsB_EG + AnsB_BG, (iii) AnsB_CBH + AnsB_BG, and (iv) AnsB_EG + AnsB_CBH + AnsB_BG, following a previously reported protocol^32^.

### Screening of multi-enzyme co-secretion strains

Plasmids encoding individual enzymes and their combinations were transformed into *E. coli* BL21 (DE3) via the heat shock method. Recombinant strains were cultured in LB medium supplemented with either 100 μg/mL ampicillin or 25 μg/mL chloramphenicol. Protein expression was induced with 0.1 mM IPTG at OD600 = 0.6, followed by incubation at 30 °C for 2, 4, 6, and 8 hours. Post induction, cultures were centrifuged (8000 ×g for 10 min), and supernatants were sterile filtered through a 0.22 μm membrane to collect secreted enzymes. For AnsB-tagged EG, CBH, and their combinations, 50 μL of extracellular media was incubated with 1 % CMC in 50 μL McIlvaine buffer, pH 6.0, yielding a 150 μL reaction volume. Reactions were performed at 55 °C for 30 minutes at 1000 rpm in a thermomixer (Eppendorf, Hamburg, Germany). After centrifugation at 8000 rpm for 2 minutes, reducing sugars were quantified using the DNS assay by measuring absorbance at 540 nm, following a previously reported protocol. ^33^

For AnsB-tagged BG or enzyme combinations containing BG, 50 μL of extracellular media was incubated with 20 mM *p*NPGlc in McIlvaine buffer, pH 6.0, to a final volume of 100 μL at 74 °C for 5 minutes at 1000 rpm in a thermomixer. The reaction was stopped by adding 0.4 M glycine solution (pH 11), and the released *p*-nitrophenol (*p*NP) was quantified by measuring absorbance at 405 nm using a standard curve to determine product concentration.

### Verification of AnsB-mediated secretion by immunoblotting

To verify the extracellular secretion of AnsB-tagged enzymes, 5 μL of supernatant was collected from cultures at different time points (2, 4, 6, and 8 hours post-induction) and analyzed via SDS-PAGE. The separated proteins were then transferred to a PVDF membrane for Western blotting. The His-tagged AnsB fusion proteins were detected using an anti-His primary antibody, while cell lysis was monitored using an anti-MBP primary antibody since MBP (Maltose Binding Protein), a periplasmic protein in *E. coli*, serves as a lysis marker. HRP-conjugated secondary antibodies enabled chemiluminescent detection. Control experiments were performed using supernatants from cultures expressing non AnsB-tagged enzymes to evaluate the impact of the AnsB fusion on secretion efficiency. The presence of MBP in the supernatant would indicate cell lysis rather than active secretion, helping differentiate true secretion from intracellular leakage.

### Enzymatic assay on natural cellulosic substrates

The hydrolytic activity of the secreted enzyme cocktail was tested on various cellulose substrates, including commercially available microcrystalline cellulose (Avicel). Untreated microcrystalline cellulose (MCC) derived from sugarcane bagasse, nanocrystalline cellulose (NCC), α-cellulose, and untreated cellulose were a gift from Prof. S. Venkata Mohan (CSIR-IICT, Hyderabad, India). For each assay, 5 % (w/v) substrate, 100 µL of secreted culture supernatant, was mixed with 100 µL of McIlvaine buffer (pH 6.0) to a final volume of 200 µL and incubated at 55 °C for different time points. The amount of reducing sugar released was quantified using the DNS assay.

### Enzymatic assay in seawater

Seawater samples were collected from two distinct locations along Puri Beach, Odisha, India. The seawater was filtered through a 0.2 µm membrane to remove particulate matter and used without any further chemical treatment. The filtered seawater exhibited a native pH of 7.65. The enzymatic activity of the secreted AnsB-tagged cellulase cocktail was evaluated in seawater. Reactions were set up as described in the DNS assay protocol and compared against reactions conducted in McIlvaine buffer (pH 6.0), the optimal buffer for the engineered cocktail. For benchmarking, the commercial cellulase cocktail, Cellic CTec2 (Novozymes, Franklinton, NC, USA), was also evaluated in both seawater and its optimal buffer (McIlvaine buffer, pH 5.0). Reducing sugar release was measured via the DNS method to determine seawater’s impact on enzyme efficiency.

### Statistical Analysis

All experiments were conducted in triplicate unless stated otherwise, and data presented as mean ± standard deviation.

## Results and discussion

### Cloning and plasmid construction

An efficient cellulase cocktail requires a glucose-tolerant BG to prevent product inhibition, a processive CBH to break down cellulose into cellobiose from non-reducing ends, and an EG to randomly cleave internal β-1,4-glycosidic bonds in cellulose and increase substrate accessibility^34-36^. For optimal performance, all enzymes should have compatible temperature Topt and pHopt ranges. In this study, we used an engineered glucose-tolerant BG, B8CYA8tm (V169C/E173L/I246A B8CYA8) with high specific activity and long half-life^23^. Additionally, we used a CBH, G0SAE6 from *C. thermophilum ^37^* and H1AD14, a processive GH9 EG from *B. licheniformis^36^*. The GH9 family EG displayed a temperature optimum of 55 °C and pHopt 5.5-6.5 and showed broad substrate specificity and high activity, making it a suitable candidate for cellulase cocktails^36^. In 1990, Jennings et al. reported that *E. coli* L-asparaginase II (AnsB) functions as a secretory protein, potentially due to a 22-amino acid signal peptide^38^. However, this signal peptide has not yet been systematically characterized as an effective secretion tag^24^. Therefore, we sought to evaluate its secretion efficiency and explore its potential use in our engineered enzyme cocktail.

The open reading frames (ORFs) of the selected enzymes were cloned into the PBP-ORF plasmid with a C-terminal His-tag, using the EcoFlex Modular Assembly system, a Golden Gate-based synthetic biology toolkit designed for standardized cloning^32^. Additionally, a 22 amino acid secretion signal sequence from *E. coli* L-asparaginase II (*AnsB*) was incorporated into the PBP-LacZ EcoFlex modular assembly plasmid^32^ (Fig. 1). SignalP 5.0 software predicted this sequence as a potential Sec pathway secretory signal peptide^39^ (SI Fig. 1). This modular cloning approach ensured precise assembly of each functional expression cassette, incorporating the T7 promoter, pET ribosome binding site (RBS), AnsB signal tag, and BBa_B0015 double terminator. It also enabled rapid screening of different enzyme combinations and secretion signal efficiency. The T7 expression system and His-tag facilitated downstream immunoblot detection, while the AnsB tag was included to promote protein secretion into the extracellular medium. This strategy simplifies enzyme activity assays and potentially reduces downstream purification costs.

**Figure 1.**
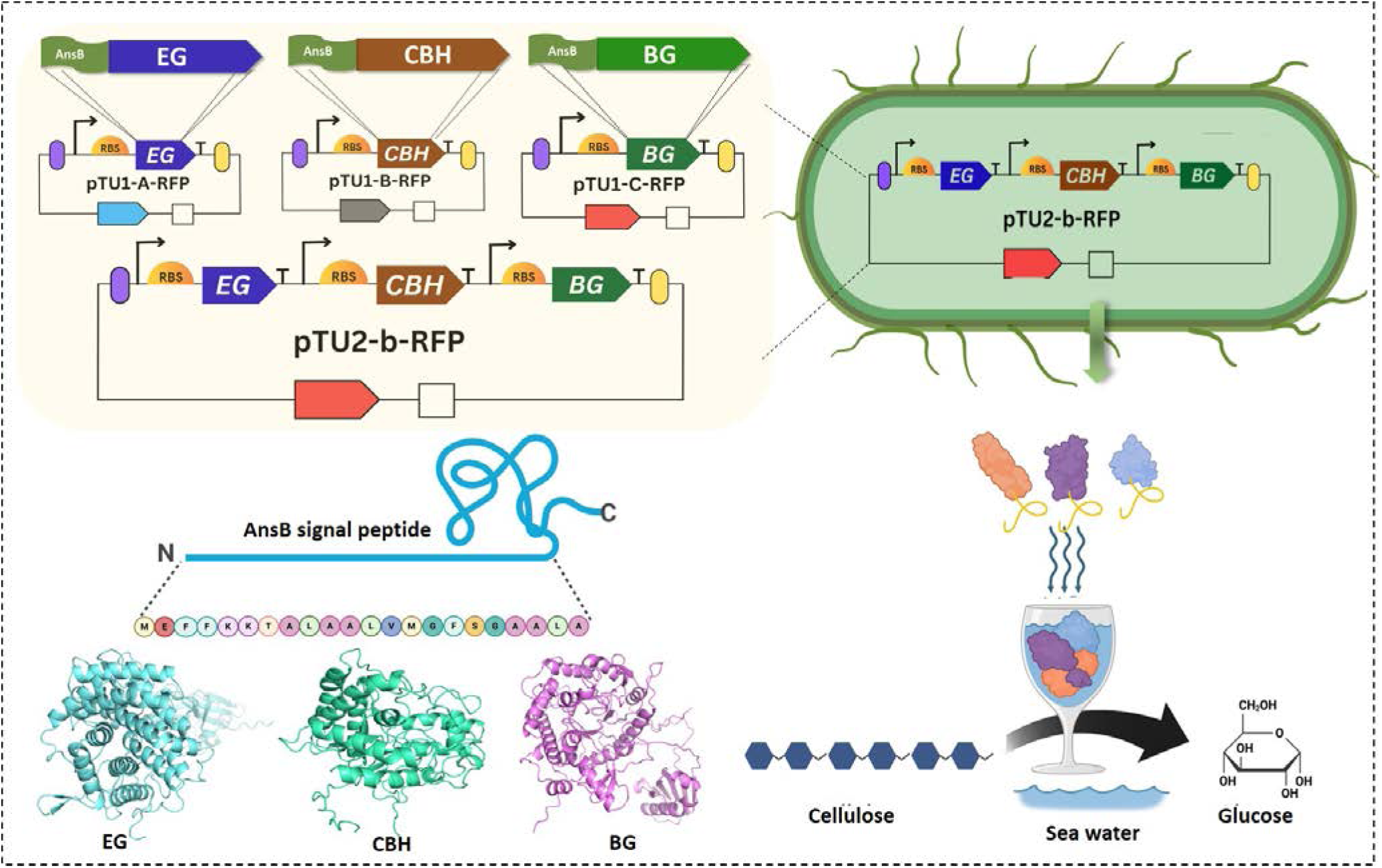
Schematic of plasmid constructs enabling extracellular secretion of a cellulase enzyme cocktail in *E. coli.* (Top left) Individual plasmids express endoglucanase (EG), cellobiohydrolase (CBH), and β-glucosidase (BG), each fused to an N-terminal AnsB signal peptide. These were combined into a single construct (pTU2-b-RFP) for co-expression. (Top right) Engineered E. coli secretes the enzyme cocktail. (Middle) Sequence and cartoon representing the AnsB signal peptide. (Bottom) Structures of EG, CBH, and BG. (Bottom right) Secreted enzymes degrade cellulose in seawater to release glucose.

### Immunoblotting and secretion analysis

To assess the secretion efficiency of AnsB-tagged cellulases, *E. coli* BL21(DE3) cultures were grown in shake flasks at 37 °C. Protein expression was induced with 1 mM IPTG at an OD600 of 0.6, followed by a temperature lowering to 30°C. Samples were collected at 2, 4, 6, and 8 hours post-induction for extracellular enzymatic activity assays. Parallel cultures expressing cellulases without the AnsB secretion tag served as controls for comparison. The enzymatic activity was measured using CMC (for EG and CBH) and *p*NPGlc (for BG) as substrates, with the reaction products quantified via the DNS assay and *p*NPGlc assay. AnsB-tagged EG, CBH, and BG exhibited 35 %, 90 %, and 72 % higher than that of non-tagged EG, CBH, and BG (Fig. 2e-g). The relative activity values presented at different time points were normalized to the 8-hour post-induction levels of each tagged enzyme to enable time-course comparisons of secretion efficiency. The culture supernatants from AnsB-tagged enzymes exhibited significantly higher enzymatic activity compared to their non-tagged counterparts, and confirmed that the AnsB tag enhances enzyme secretion into the culture medium, likely via the Sec secretion pathway.

**Figure 2.**
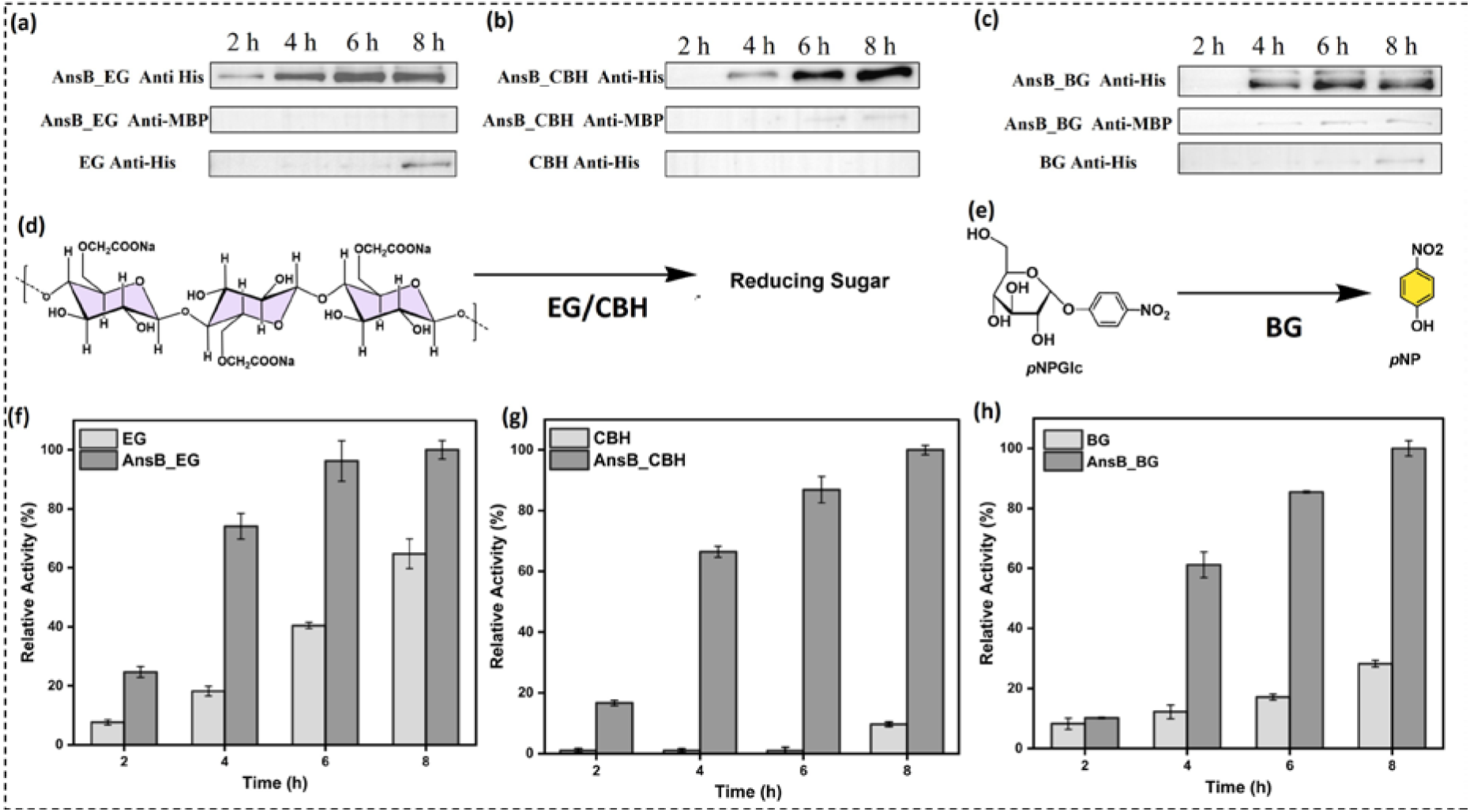
Western blot analysis of the supernatant collected at 2,4,6 and 8 hours post-induction for (a) EG and AnsB tagged EG, (b) CBH, and AnsB tagged CBH, and (c) BG and AnsB tagged BG. An anti-His antibody was used to detect His tagged secreted enzymes in the culture medium to confirm the presence of AnsB-tagged proteins in the extracellular fraction. (d) Substrate hydrolysis mechanisms of secreted cellulases. (d) Schematic of endoglucanase (EG) and cellobiohydrolase (CBH) activity on carboxymethyl cellulose (CMC), leading to the release of reducing sugars. (e) β-Glucosidase (BG) catalyses hydrolysis of p-nitrophenyl-β-D-glucopyranoside (pNPGlc) to release p-nitrophenol (pNP), enabling colorimetric activity detection. (e) Schematic representation of the catalytic performance of BG and AnsB-tagged BG. Comparative enzymatic activity of secreted media: (f) EG with and without AnsB tag, (g) CBH with and without AnsB tag, and (h) BG with and without AnsB tag. Data represent mean ± standard deviation (n = 3).

Western blot analysis was performed to rule out cell lysis as a source of extracellular enzymes activity. Supernatants from different time points were analyzed via SDS-PAGE and immunoblotting. The His-tagged cellulases in the supernatant were detected using an anti-His antibody, while an anti-MBP antibody assessed periplasmic leakage. The anti-His western blots revealed prominent bands for AnsB-tagged enzymes in the extracellular media at 2, 4, 6, and 8 hours, confirming active secretion (Fig. 2a-c). In contrast, MBP was undetectable at 2 and 4 hours, indicating minimal or no cell lysis. Faint MBP bands at 6 and 8 hours suggested minor cell lysis^14^, likely due to prolonged culture conditions and enzyme overexpression (Fig. 2a-c and SI Fig. 2). Untagged controls expressing non-tagged cellulases showed no detectable His-tagged proteins early on, further supporting secretion specificity.

Several previous studies highlight the variability on secretion efficiency of heterologous proteins in *E. coli* depending on signal peptides and cultivation methods. Typically, the efficiency and yield of secreted proteins are strongly influenced by the specific properties of the protein of interest, including its length, folding kinetics, and dependence on the Sec secretion pathway^40^. For instance, recombinant L-asparaginase II tagged with the pelB signal sequence achieved titers of approximately 110 mg/L in shake flask conditions^28^, while staphylokinase with the OmpA signal sequence yielded only 0.005 g/L under similar conditions^25^. The PhoA tag has been used to express murine endostatin at 40 mg/L^41^, and OmpA has also facilitated the secretion of human growth hormone at ∼76 mg/L^42^ and PNGase F at 8 mg/L^26^. Notably, higher titers were achieved in fed-batch processes, for example, 5.2 g/L of alkaline phosphatase containing the Bacillus sp. endoxylanase signal sequence^27^ and 2 g/L of levan fructotransferase using a lacZ-derived signal peptide^43^. These results underscore the critical roles of signal peptide selection and cultivation strategy in secretion efficiency.

In comparison to other secretion systems, our AnsB-derived signal peptide achieved titers of 0.2 to 0.4 g/L (EG 0.24 g/L, CBH 0.33 g/L and 0.4 g/L) in shake-flask cultures, as quantified by immunoblotting (SI Fig. 3). These yields are comparable or superior to many previously reported systems under similar conditions (SI Table 3), demonstrating the efficacy of the AnsB tag for cellulase secretion. The secretion system offers key practical benefits. It eliminates the need for cell disruption, intracellular extraction, or solubilization of inclusion bodies. It enables the direct use of culture supernatants in biomass hydrolysis, streamlining downstream processing. Furthermore, it facilitates high-throughput screening of enzyme libraries in extracellular media, supporting directed evolution and optimization without cell lysis. Since secretion remains a major bottleneck in recombinant protein production in *E. coli*, especially for industrially relevant hydrolases, our work establishes the AnsB signal peptide as a versatile and scalable tool. It provides a proof-of-concept for leveraging native or engineered signal peptides to enhance secretion efficiency, opening new possibilities for bioprocess optimization and enzyme engineering.

### Synergistic effects of AnsB-tagged enzyme combinations

After confirming individual enzyme secretion, we constructed AnsB-tagged cellulase combinations to investigate their synergistic effects. Four enzyme combinations were designed: (i) AnsB_EG + AnsB_CBH, (ii) AnsB_EG + AnsB_BG, (iii) AnsB_CBH + AnsB_BG, and (iv) AnsB_EG + AnsB_CBH + AnsB_BG. Western blot analysis was performed to validate the extracellular secretion of the enzyme combinations. Shake flask cultures of *E. coli* BL21(DE3) expressing individual and combined enzyme constructs were grown in LB medium to an OD600 of 0.6, followed by induction with 1 mM IPTG. The cultures were incubated for 8 hours at 30°C before harvesting. Cells were pelleted by centrifugation at 8000 rpm for 10 minutes, and 5 µL of the collected culture supernatant was subjected to SDS-PAGE and western blot. Western blot detection confirmed extracellular secretion for all combinations (Fig. 3a), indicating that the AnsB secretion tag remains effective even when multiple tagged enzymes are co-expressed.

**Figure 3.**
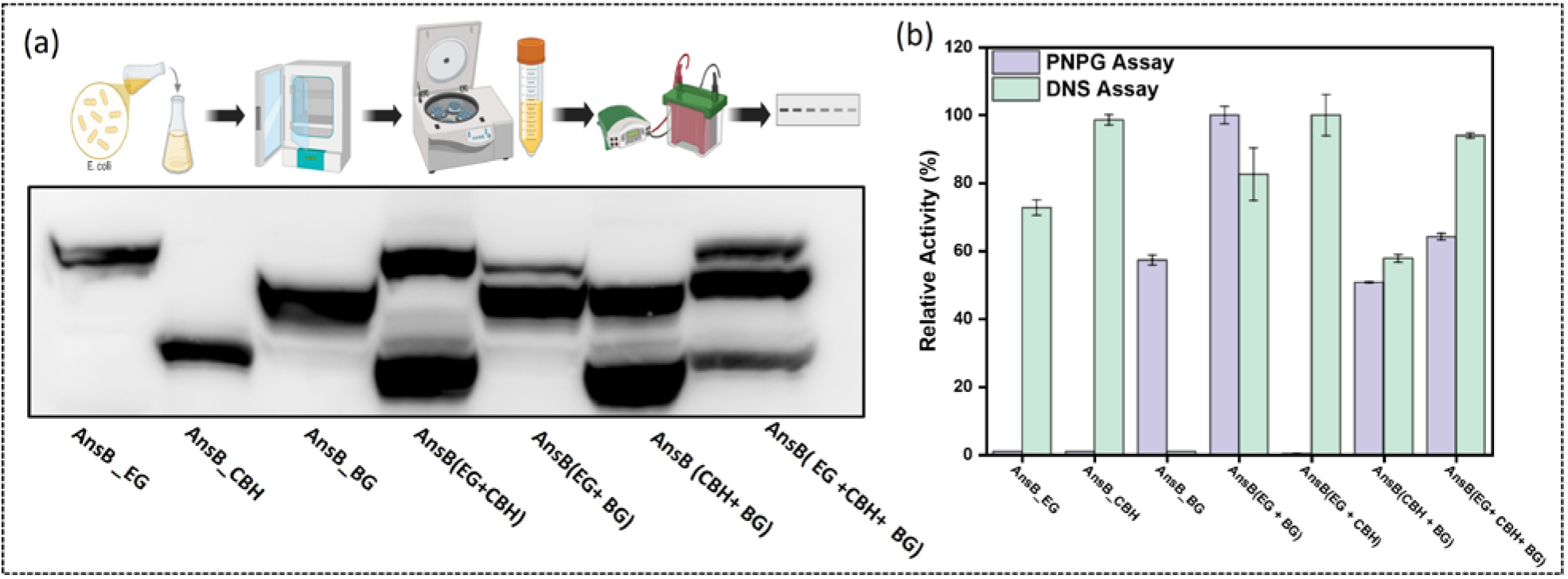
(a) Western blot analysis of AnsB-tagged secreted enzymes in culture supernatant using an anti-His antibody. To confirm successful secretion. The analyzed samples include EG, CBH, BG, EG + CBH, EG + BG,CBH + BG and EG + CBH + BG. (b) Enzymatic activity of secreted AnsB-tagged enzymes (EG, CBH, BG, EG + BG, EG + CBH, CBH + BG and EG + CBH + BG) was assessed by pNPGlc and DNS assay. pNPGlc assay quantifies β-glucosidase activity, while DNS measures reducing sugar release. Data represent mean ± standard deviation (n = 3).

The enzymatic activity of individual enzymes and their combinations was evaluated using CMC and *p*NPGlc as substrates to validate the constructed enzyme combinations further and assess their synergistic effects. For both assays, the relative activity of the combination showing the highest activity was defined as 100 %, and all other values were normalized accordingly. The AnsB-tagged EG, CBH, BG alone showed 72 %, 98 % and 0 % relative activity, respectively, while their combinations, EG + CBH, EG + BG, CBH + BG, and EG + CBH + BG, exhibited 82 %, 100 %, 56 %, and 94 % relative activity, respectively. As expected, AnsB-tagged BG alone did not hydrolyse CMC, as BG primarily acts on di-oligosaccharides rather than polysaccharides (Fig. 3b). To confirm the secretion and activity of BG in the combinations, *p*NPGlc hydrolysis was assessed by measuring *p*-nitrophenol (*p*NP) release. The AnsB-tagged EG, CBH, and EG+ CBH had 0 % activity (no *p*NP release) while BG alone showed 100 % activity. Its combinations with EG + BG, CBH + BG, and EG + CBH + BG, showed 58 %, 50 %, and 64 % relative activity, respectively (Fig. 3b). These results confirm that the AnsB-tagged EG+ CBH+ BG cocktail efficiently hydrolyzes both polymeric (CMC) and oligomeric (*p*NPGlc) substrates, demonstrating functional synergy. Enhanced activity in mixed combinations compared to individual enzymes suggests sequential action. EG/CBH initiate cellulose depolymerization, followed by monomer release by BG, may occur efficiently even in a mixed extracellular environment. This supports the design of a modular secretion system where each component retains its function without mutual interference.or degradation. Based on this comprehensive activity profile, we selected the AnsB tagged EG + CBH + BG combination for further characterization.

Previous studies by Zhang et al. reported that the YebF protein of *E. coli* facilitates extracellular secretion when fused to target proteins^14^. To benchmark performance, we fused YebF to our enzyme cocktail (EG + CBH + BG) using modular assembly and compared its secretion efficiency with the AnsB-tagged cocktail. After 8 hours of induction, supernatants were collected and analysed using DNS and *p*NPGlc assays. The AnsB-tagged enzymes outperformed the YebF-tagged ones by yielding 5-fold higher reducing sugars by DNS assay and an 8-fold higher *p*NP production by the *p*NPGlc assay (SI Fig. 4). These results highlight the AnsB-tagged enzyme cocktail’s superior secretion efficiency than the YebF-tagged counterpart. Unlike YebF, which may be more suited for small or single-domain proteins, the AnsB tag accommodates diverse and multi-domain hydrolases with higher secretion efficiency. Future work could explore combining AnsB with signal amplification systems or co-expression with membrane-permeabilizing factors^44^ to further optimize industrial applications.

### Optimization of Growth Conditions for Enhanced Enzyme Secretion

To maximize the secretion of the AnsB-tagged EG + CBH + BG enzyme cocktail, we optimized key growth parameters, including temperature, culture media, induction timing, and inducer concentration, similar to what was done earlier. The highest reducing sugar release from CMC, measured via the DNS assay, occurred when induction was carried out at OD₆₀₀ = 0.9. In contrast, the *p*NPGlc assay (measuring β-glucosidase activity) showed peak *p*NP conversion at OD₆₀₀ = 0.6, though activity at OD₆₀₀ = 0.9 remained nearly similar (96 % compared to activity at OD₆₀₀ = 0.6). Given the consistently high enzymatic activity across both assays, OD₆₀₀ = 0.9 was selected as the optimal induction point (Fig. 4a). Inducing at this mid to late exponential phase likely balances optimal cell metabolic activity with sufficient biomass for high-level protein expression and secretion. To optimize the ideal inducer concentration, we tested IPTG concentrations ranging from 0.1 to 10 mM and evaluated enzymatic activity after 8 hours. The highest activity was observed at 0.1 mM IPTG, suggesting that lower IPTG concentrations favor efficient secretion, while higher concentrations may impose metabolic stress. Therefore, 0.1 mM IPTG was chosen for further optimization (Fig. 4b). This finding aligns with previous findings where excessive overexpression can saturate the secretion machinery and reduce yield due to cellular toxicity^45^.

**Figure 4.**
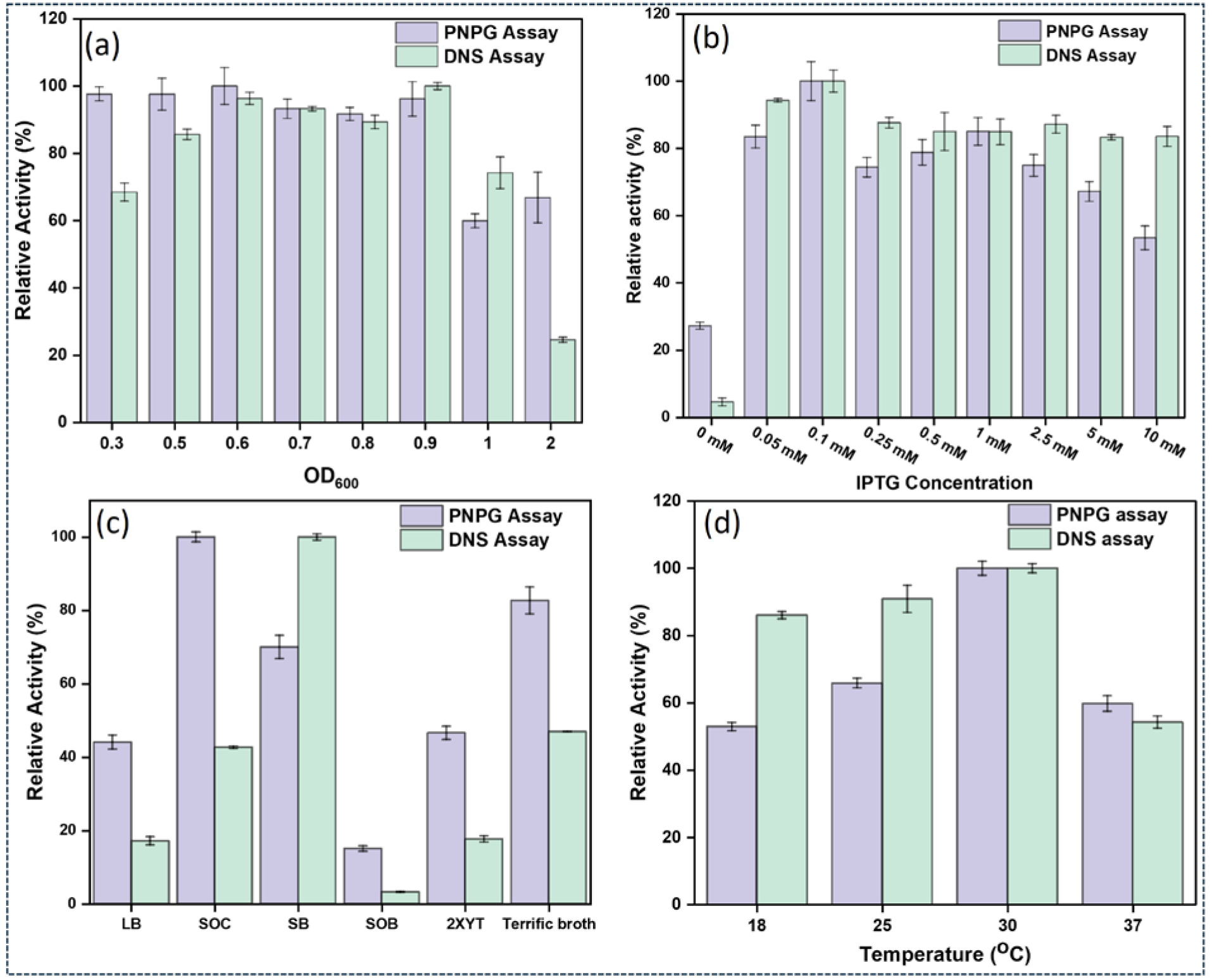
Optimization of secretory enzymatic cocktail culture. (a) Catalytic activity of AnsB-tagged cocktail secreted media at different OD₆₀₀ values ranging from 0.3 to 2 upon IPTG induction. (b) Catalytic activity of AnsB-tagged cocktail secreted media at varying IPTG concentrations (0 mM to 10 mM). (c) Catalytic activity of AnsB-tagged cocktail secreted media in different nutrient media. (d) Catalytic activity of AnsB-tagged cocktail secreted media at different growth temperatures ranging from 18 to 37 °C. Activities were measured via DNS and *p*NPGlc assay and normalized to 100 % under optimum conditions. Data represent mean ± standard deviation (n = 3).

Since media optimization is a crucial step for industrial scalability, six different media, LB, SOB, YT, Terrific, SB, and SOC, were tested by culturing *E. coli* at 30 °C. After 8 hours of growth, cells were harvested by centrifugation at 8000 rpm for 10 minutes, and the supernatant was used for activity assays. SOC medium outperformed others, yielding 5-fold higher reducing sugar production than LB (Fig. 4c). A control experiment with an empty plasmid confirmed that the observed activity was from secreted enzymes, not media-derived sugars. The superior performance of SOC medium may be attributed to its rich nutrient profile, which enhances cell growth and secretion efficiency.

To determine the optimal growth temperature for enzyme secretion, *E. coli* cultures expressing the AnsB-tagged EG + CBH + BG enzyme cocktail were induced with 0.1 mM IPTG and incubated at 18 °C, 25 °C, 30 °C, and 37 °C. The highest activity was observed at 30 °C, likely due to an optimal balance between protein folding and secretion efficiency (Fig. 4d). Higher temperatures (e.g., 37 °C) may increase protein misfolding or proteolysis, while lower temperatures slow growth and expression rates. By integrating these optimized parameters, including OD₆₀₀ at induction, IPTG concentration, culture medium, and temperature, we increased the secretion of the AnsB-tagged cellulase cocktail from 0.31 g/L to 0.74 g/L (SI Fig. 5), demonstrating a significant enhancement over unoptimized conditions.

### Enzymatic properties of AnsB-Tagged EG + CBH + BG cocktail

To determine the optimal conditions for the AnsB-tagged EG + CBH + BG enzyme cocktail, we assessed its activity across a pH range of 4.0 to 8.0 using McIlvaine buffer. Enzymatic activity was measured via the DNS assay with carboxymethyl cellulose (CMC) as the substrate. The cocktail exhibited peak relative activity at pH 6.0 (Fig. 5a), confirming this as the optimal pH for synergistic hydrolysis. Notably, the cocktail retained more than 80% of its maximal activity across a broad pH range of 4.5 to 8.0, demonstrating robust pH stability and operational flexibility for diverse bioprocessing applications. This wide pH tolerance enhances its suitability for industrial processes where pH fluctuations are common.

**Figure 5.**
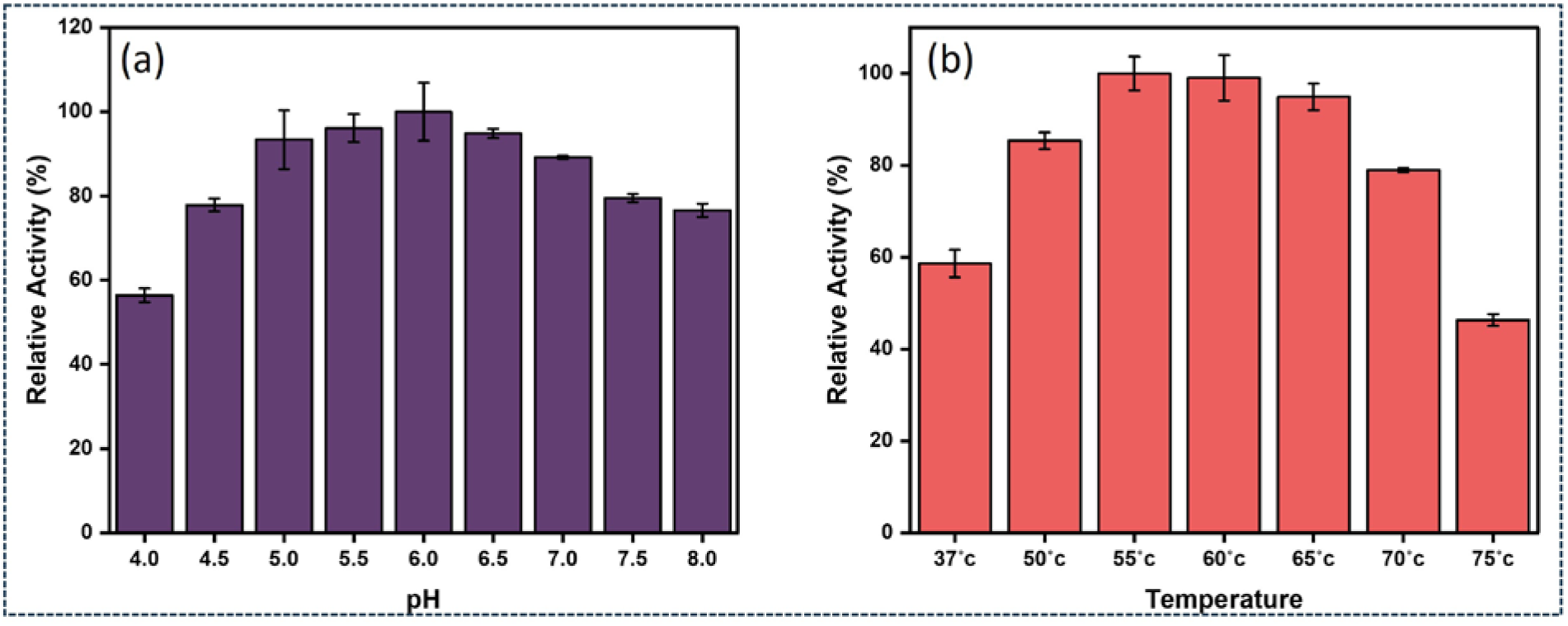
Optimization of pH and temperature for secretory enzymatic cocktail culture. (a) Catalytic activity of AnsB-tagged cocktail secreted media at different pH values. (b) Catalytic activity of AnsB-tagged cocktail secreted media at different temperatures. Activities were measured via DNS assay and normalized to 100 % under optimum pH and temperature conditions. Data represent mean ± standard deviation (n = 3).

We next evaluated thermal stability by testing the enzyme cocktail at temperatures ranging from 37°C to 75°C. Maximum activity was observed at both 55°C and 60°C, with negligible differences between these temperatures (Fig. 5b), indicating strong thermal stability and catalytic efficiency. This property is particularly advantageous for biomass saccharification, where higher temperatures improve reaction kinetics and reduce contamination risks. Additionally, the enzyme cocktail displayed excellent storage stability when stored at 4 °C for one month, without any detectable loss in activity. This long-term stability highlights its industrial potential, ensuring reliable performance during transport or between processing cycles.

### Hydrolysis of natural substrates by AnsB-Tagged EG + CBH + BG cocktail

We evaluated the enzymatic activity of our AnsB-tagged enzyme cocktail on various cellulose-based substrates, including commercially available microcrystalline cellulose (Avicel), untreated microcrystalline cellulose (MCC), nanocrystalline cellulose (NCC), α-cellulose, and untreated cellulose.

The enzyme cocktail effectively hydrolysed all substrates, producing reducing sugar concentrations of 1.97, 1.57, 1.48, 1.29, and 0.63 g/L, respectively. Avicel yielded the highest sugar release, likely due to its reduced crystallinity and enhanced enzyme accessibility. MCC and NCC also showed relatively high hydrolysis, suggesting that their nanoscale dimensions, reduced fiber packing, and microstructural refinement facilitate efficient enzyme-substrate interactions^46-48^. In contrast, α-cellulose and untreated cellulose showed lower yields, likely due to their higher crystallinity and dense fibrous structure, hindering enzyme penetration and catalytic efficiency^49, 50^. These results highlight the effectiveness of the AnsB-tagged cocktail in saccharifying diverse lignocellulosic materials and emphasize the critical role of substrate pretreatment in optimizing bioconversion efficiency.

### Effect of seawater on enzyme cocktail activity

To date, the majority of cellulose saccharification processes for biofuel production rely on freshwater as the primary reaction medium. Studies have shown that the water footprint for producing one litre of bioethanol can range from 790 L (sugar beet, France) to over 11,030 L (molasses ethanol, Thailand), depending on feedstock and location^29-31^. With the growing global freshwater crisis, identifying sustainable and abundant alternatives is critical. Seawater, abundant in coastal regions and cost-effective, offers a promising solution. However, its feasibility depends on enzyme halotolerance and catalytic efficiency under high-salinity conditions.

To address this, we evaluated seawater as a sustainable medium for cellulose hydrolysis. Remarkably, the AnsB-tagged enzyme cocktail retained nearly 100 % saccharification activity in unadjusted seawater, matching its performance in optimal cocktail buffer conditions (Fig. 7a). To our knowledge, this is among the first reports of a cellulase cocktail functioning effectively in seawater. In contrast, the commercially available Cellic CTec2 cocktail exhibited a ∼75 % activity reduction in seawater compared to its activity in its optimal buffer (Fig. 7b). While seawater’s high salinity and alkalinity typically inhibit cellulase activity, our engineered cocktail maintained full hydrolytic activity, offering an eco-friendly and resource-efficient approach to biomass conversion. These findings provide strong proof of concept for future *E. coli* strain engineering. Additionally, our genetic modifications and secretion strategies are modular and adaptable, making them suitable for integration into other industrial microbial hosts, expanding their biotechnological applications. Future work could focus on elucidating the structural basis of enzyme halotolerance, enabling the rational design of seawater-compatible biocatalysts for use in marine-integrated biorefineries.

**Figure 6.**
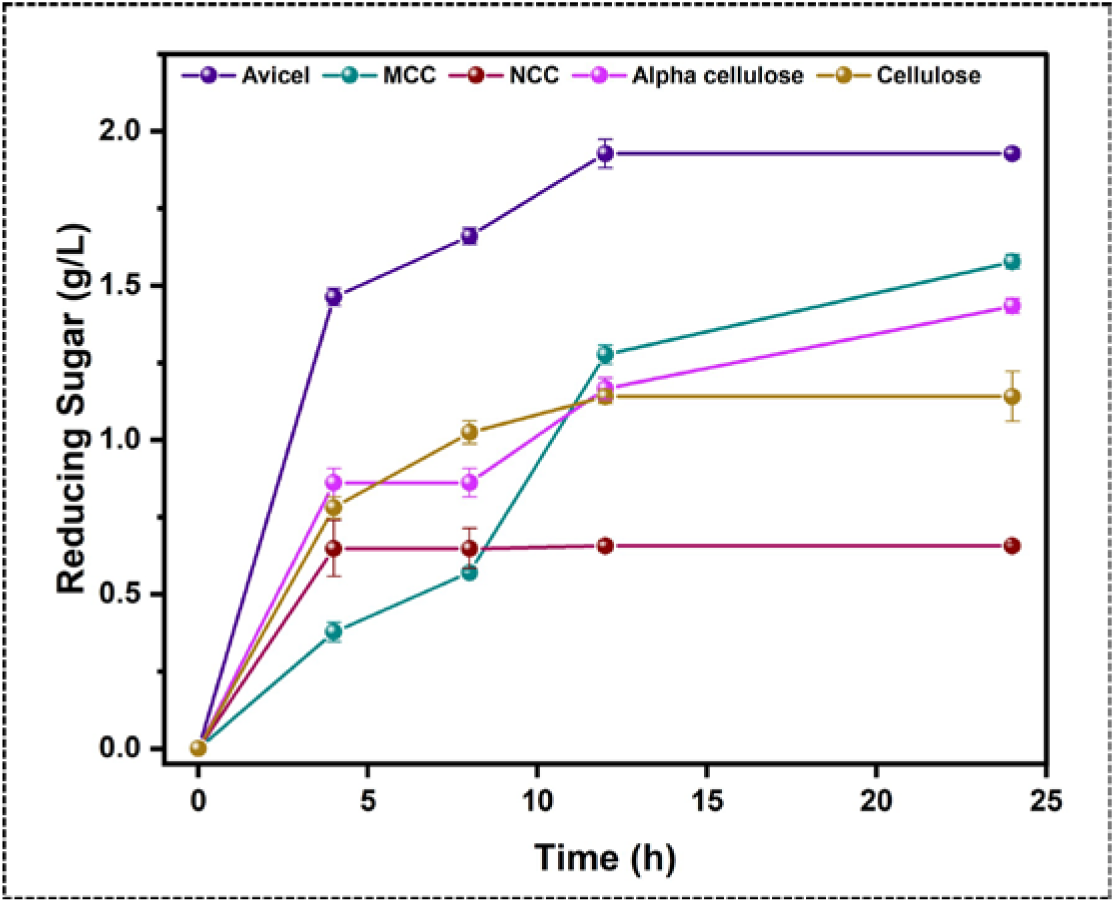
Time-course of reducing sugar release (g/L) from various natural cellulosic substrates upon hydrolysis by the AnsB-tagged cellulase cocktail. The catalytic performance was evaluated using Avicel (microcrystalline cellulose), MCC (microcrystalline cellulose), NCC (nanocrystalline cellulose), alpha-cellulose, and cellulose as substrates. The enzyme cocktail showed the highest hydrolytic efficiency on Avicel, followed by MCC and NCC, indicating its robust activity on crystalline cellulose forms. Data represent mean ± standard deviation (n = 3).

**Figure 7.**
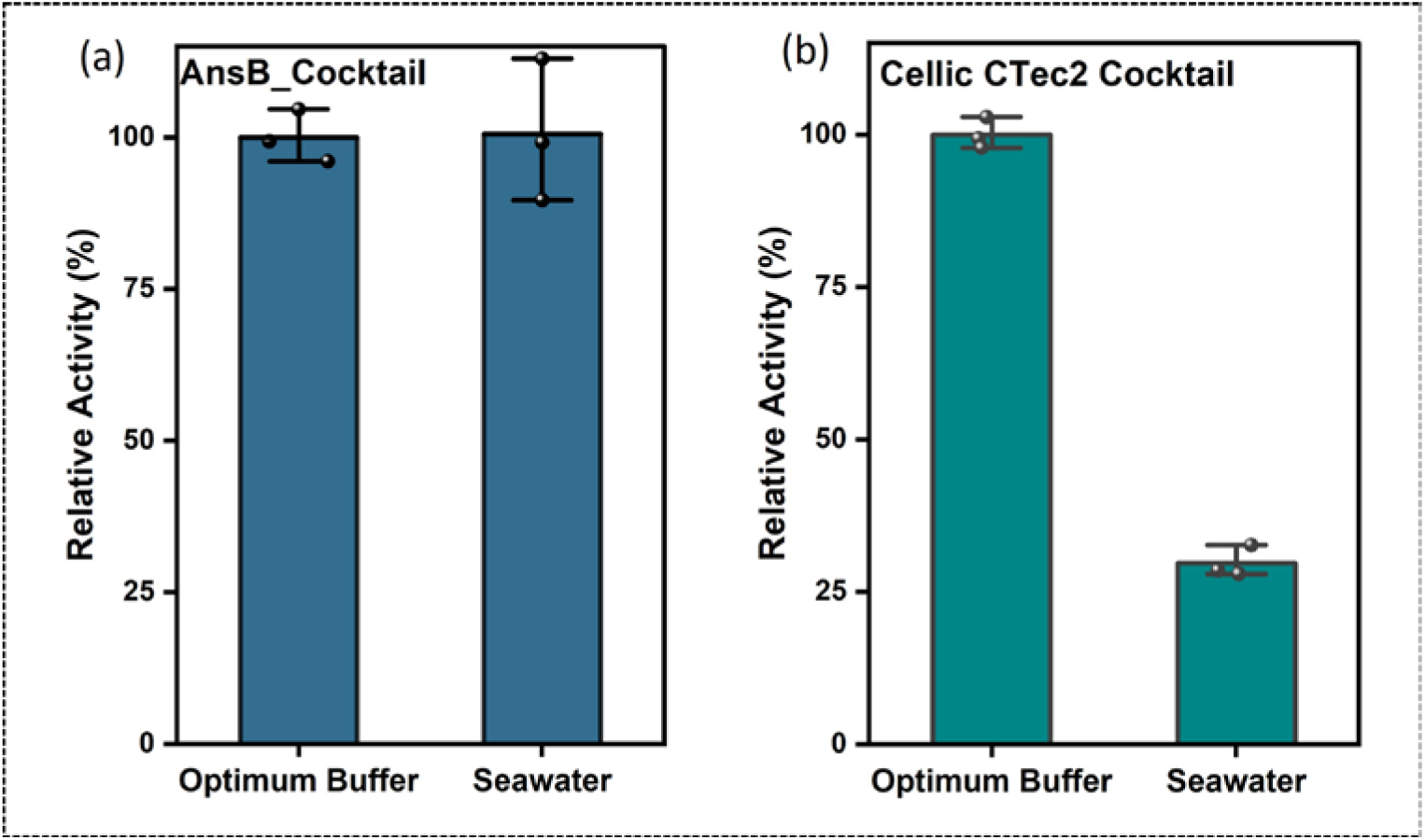
Comparison of relative specific activities of Ans-B tagged cellulase cocktail in its optimum buffer and natural sea water compared to Cellic Ctec 2 (Novozyme, Bangalore India) in its optimum buffer and in seawater. (a) AnsB-tagged cellulase cocktail retained similar activity in seawater, with no significant loss compared to reaction in its optimum buffer. (b) Commercial Cellic CTec2 Cocktail showed a substantial reduction in activity in seawater relative to activity in its optimum buffer. Activities were measured via DNS assay and normalized to 100 % under optimum buffer conditions. Data represent mean ± standard deviation (n = 3).

### Conclusions

We successfully engineered an AnsB-tagged cellulase cocktail (EG + CBH + BG) in *E. coli*, achieving efficient extracellular secretion and robust enzymatic activity. The cocktail demonstrated the synergistic hydrolysis of diverse cellulose substrates, maintained high activity in seawater, and showed excellent stability across a broad pH and temperature range. By optimizing expression parameters, we further enhanced secretion yields. Compared to alternatives like YebF, the AnsB provided superior secretion efficiency. These results highlight the potential of our enzyme system for sustainable biomass saccharification, particularly in saline and resource-limited environments, advancing the development of industrially viable biofuel production processes.

## ASSOCIATED CONTENT

### Author Contributions

S.D. supervised the project. M.G., S.D., conceived and designed the project. M.G. designed all the experiments, cloned and assembled AnsB tagged constructs performed assay, western blots, collected and analysed the characterization data, and prepared all the figures. A.S. performed cloning, secretion study optimization, western blots, and enzymatic assays. Suksham cloned yebF tagged enzymes and performed enzymatic assay experiments. D.C. helped in initial cloning and experiments. The manuscript was written through the contributions of M.G. and S.D. All authors have given approval to the final version of the manuscript.

### Notes

The authors declare no competing financial interests.

## ACKNOWLEDGMENT

MG acknowledges DST-INSPIRE for a Senior Research Fellowship. SD acknowledges SERB Core Research Grant ([CRG/2023/002111](https://serbonline.in/SERB/MenuPage)), and DBT, Government of India (grant number BT/PR47801/BCE/8/1812/2023). AS acknowledges DST-INSPIRE scholarship. We acknowledge Dr. S Venkata Mohan for providing pretreated natural cellulose substrates.

## Toc

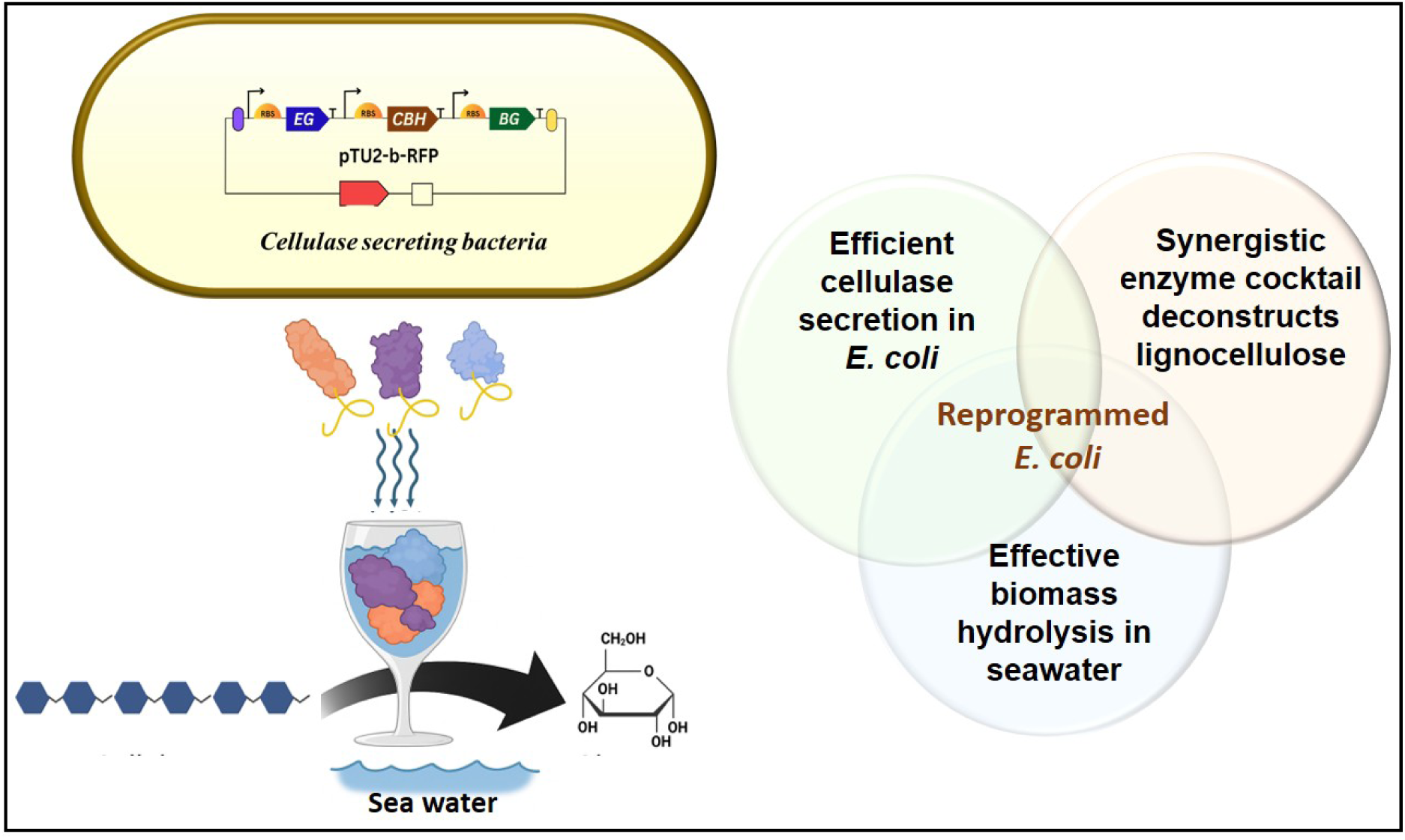

