## Supplementary Files for "Reprogrammed *E. coli* for secretion of thermophilic cellulase cocktail for seawater-compatible lignocellulosic bioprocessing"

### Reprogramming *E. coli* for secretion of thermophilic cellulase cocktail for biomass degradation in seawater

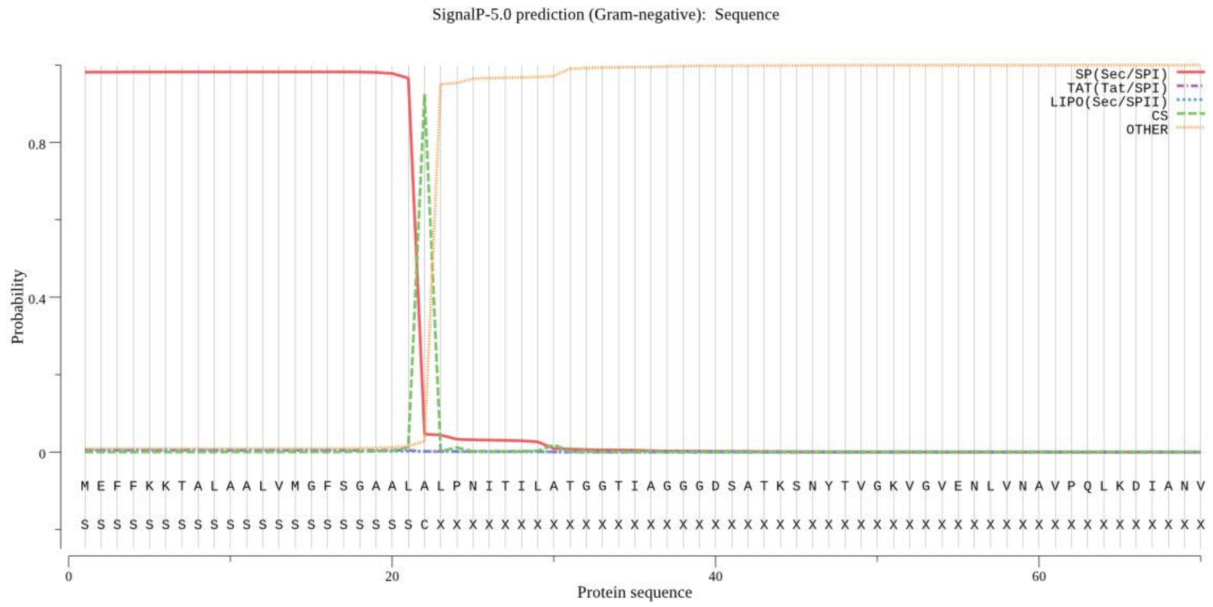


**Figure S1.** Signal peptide prediction of the AnsB N-terminal sequence using SignalP 5.0. The plot displays the predicted probability scores for different secretion pathways in gram-negative bacteria. The red line indicates a high probability for Sec/SPI-dependent secretion, confirming the presence of a cleavable signal peptide at the N-terminus. The cleavage site (CS), predicted between positions 21 and 22, is marked by a sharp drop in Sec signal probability and a peak in the CS score, supporting the potential role of the AnsB signal peptide in mediating Sec-pathway-dependent secretion.

### Western Raw Images

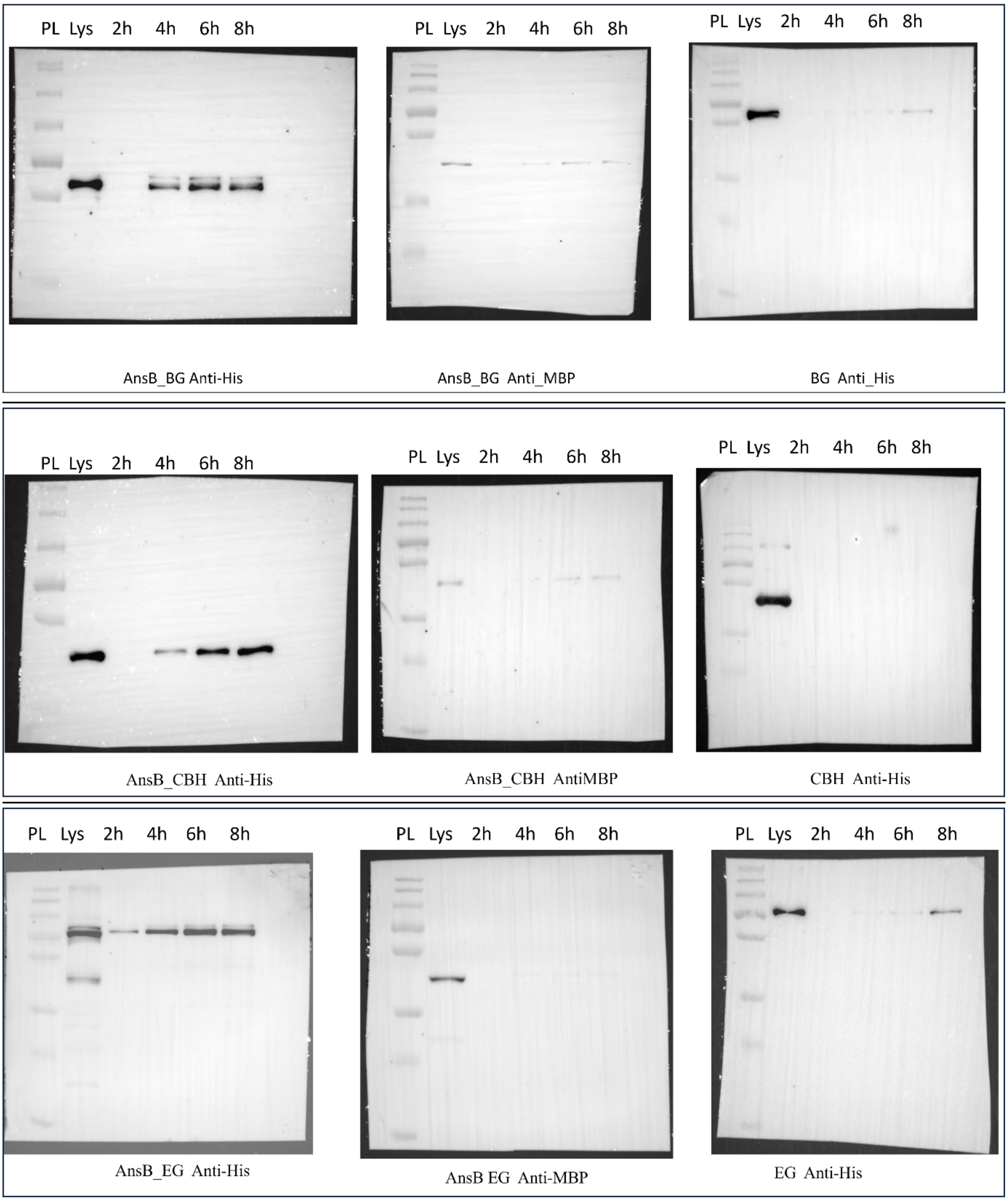


**Figure S2.** Unprocessed Western blot images corresponding to the main figure panels (Fig.2 a-c), showing the secretion profile.

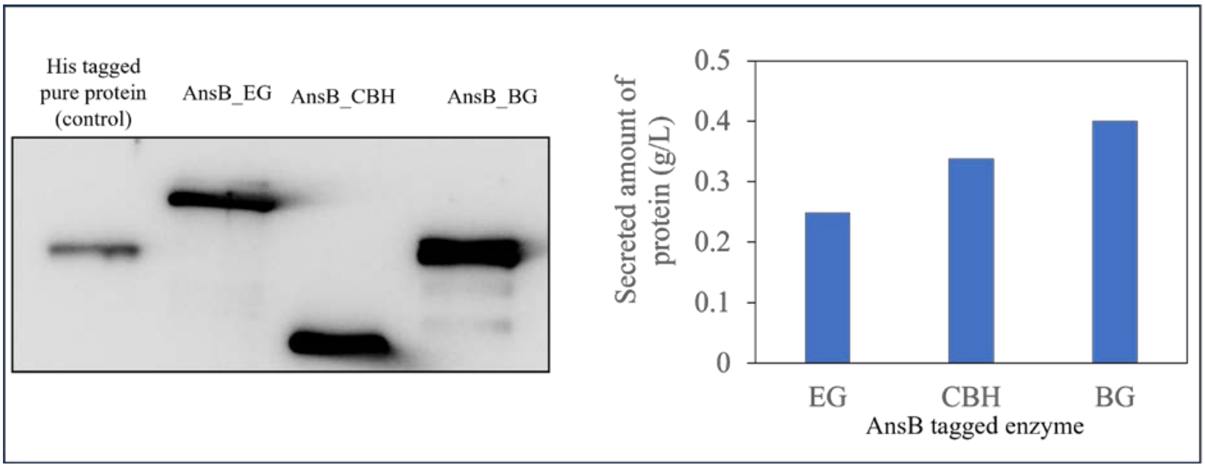


**Figure S3.** Quantification of secreted AnsB tagged EG, CBH, and BG enzymes using Western blot analysis. The left panel shows representative Western blot bands for the secreted enzymes EG, CBH, and BG from culture supernatants of *E. coli* expressing AnsB-tagged constructs. The right panel presents quantification of secreted protein levels (g/L), determined by densitometric analysis of band intensities using ImageJ software.[2]

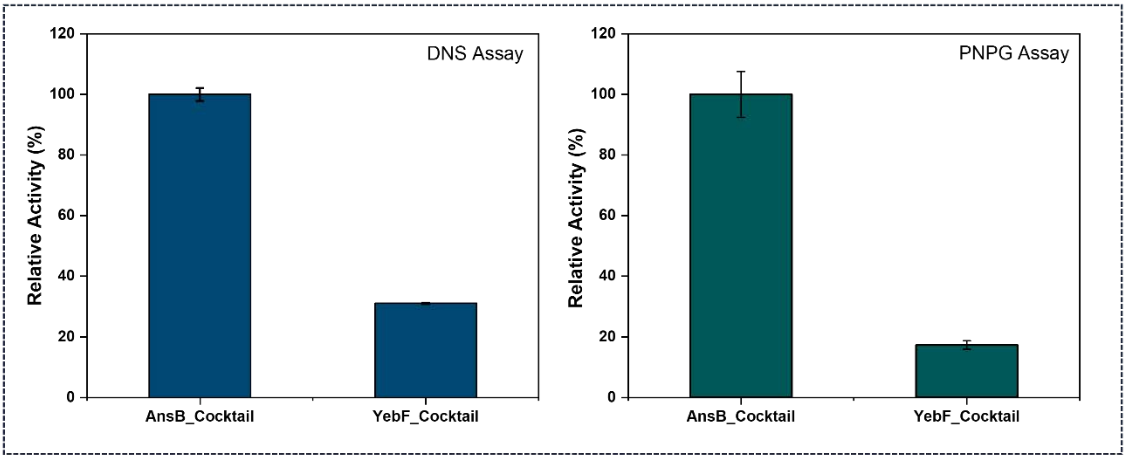


**Figure S4.** Comparison of the enzymatic activity of secretory cellulase cocktails tagged with either AnsB or YebF by DNS assay (left panel) to measure reducing sugars, or *p*NPGlc assay to detect glucose released (right panel).

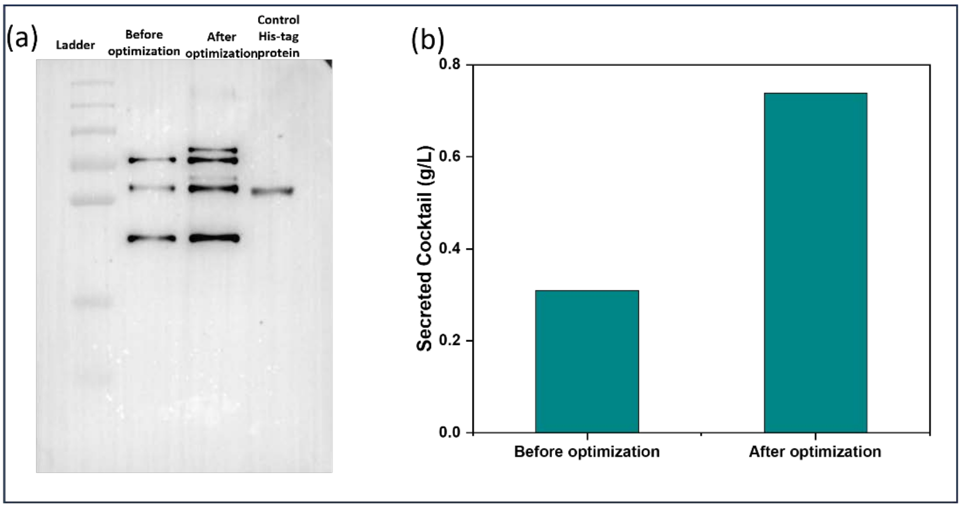


**Figure S5.** Quantification of secreted cellulase cocktail before and after optimization of growth conditions (details in methods and results). (a) Western blot analysis of secreted proteins from *E. coli* expressing the cellulase cocktail before and after secretion optimization, with the His-tagged purified enzyme cocktail used as a control. (c) Quantification of total secreted cocktail based on band intensity analysis using ImageJ software.

**Table S1.** Oligonucleotides used for cloning and engineering the cellulase genes

| Primer | DNA sequences |
|---|---|
| AnsB tag_FP | GGCAATTCCATATGGGTCTCATAAAATGGAGTTTTTCAAAAAGAC |
| AnsB tag_RP | CACATGCATGCGGTCTCTTATGTGCCAATGCTGCAC |
| EG_FP | GTATGGTCTCACATATGGCGGACATTGATTTG |
| EG_RP | GCATGCGGTCTCTTCGATTAATGATGGTGATGGTGGATGGT |
| CBH_FP | GTATGGTCTCACATATGGCGCCATCCTTCACG |
| CBH_RP | ATTAGGTCTCTTCGATCAATGGTGGTGGTGATGATG |
| BG_FP | CGCTACGGTCTCACATATGGCGAAGATTATTTTTCCAGAAGAC |
| BG_RP | ATTAGGTCTCTTCGAATGATGATGATGATGATGATTTG |

**Table S2.** Strains and plasmids used in this study

| Strains or Plasmids | Characteristics | References |
|---|---|---|
| pBP-T7 | T7 promoter | 1 |
| pBP-Tag_linker | PET RBS with tag linker | 1 |
| pBP-ORF | Empty vector to clone ORF | 1 |
| pBP-Bba_B0015 | Synthetic double terminator | 1 |
| pBP-EG | Endoglucanase at level 0 | This work |
| pBP-CBH | Cellobiohydrolase at level 0 | This work |
| pBP-BG | Beta-glucosidase at level 0 | This work |
| pBP_AnsB | AnsB tag at level zero | This work |
| pTU-1A- AnsB_EG (A) | Endoglucanase at level 1 | This work |
| pTU-1B- AnsB_CBH (B) | Cellobiohydrolase at level 1 | This work |
| pTU-1C- AnsB_BG ( C) | Beta-glucosidase at level 1 | This work |
| pTU-2a- AnsB_EG+ AnsB_CBH (D) | EG and CBH at level 2 | This work |
| pTU-2a- AnsB_EG+AnsB_BG (E) | EG and BG at level 2 | This work |
| pTU-2a- AnsB_CBH+ AnsB_BG (F) | CBH and BG at level 2 | This work |
| pTU-2b- AnsB_EG+ AnsB_CBH+ AnsB_BG (G) | EG, CBH and BG at level 2 | This work |
| *E. coli* BL21(DE3) | Protein expression host | ThermoFisher |
| *E. coli* Dh5α | Cloning strain | Novagen |
| E. coli BL21/(A) | E. coli BL21 containing plasmid A | This work |
| *E. coli* BL21/(B) | E. coli BL21 containing plasmid B | This work |
| *E. coli* BL21/(C) | E. coli BL21 containing plasmid C | This work |
| *E. coli* BL21/(D) | E. coli BL21 containing plasmid D | This work |
| *E. coli* BL21/(E) | E. coli BL21 containing plasmid E | This work |
| *E. coli* BL21/(F) | E. coli BL21 containing plasmid F | This work |
| *E. coli* BL21/(G) | E. coli BL21 containing plasmid G | This work |
| *E. coli* Dh5α /(A) | *E. coli* DH5 α containing plasmid A | This work |
| *E. coli* Dh5α /(B) | *E. coli* DH5 α containing plasmid B | This work |
| *E. coli* Dh5α /(C) | *E. coli* DH5 α containing plasmid C | This work |
| *E. coli* Dh5α /(D) | *E. coli* DH5 α containing plasmid D | This work |
| *E. coli* Dh5α /(E) | *E. coli* DH5 α containing plasmid E | This work |
| *E. coli* Dh5α /(F) | *E. coli* DH5 α containing plasmid F | This work |
| *E. coli* Dh5α /(G) | *E. coli* DH5 α containing plasmid G | This work |

**Table S3**. Recombinant protein secretion yields using different signal peptides in *E. coli* reported in the literature

| Protein | Signal Peptide/Tag | Amount (g/L) | Expression Mode | Reference |
|---|---|---|---|---|
| L-asparaginase II | pelB | 0.11 | Flask | 3 |
| Staphylokinase | ompA signal sequence | 0.005 | Flask | 4 |
| murine endostatin | phoA | 0.04 | Fed-batch | 5 |
| Human growth hormone | ompA | 0.076 | Flask | 6 |
| Alkaline phosphatase | Bacillus sp. endoxylanase signal sequence | 5.2 | Fed-batch | 7 |
| Levan fructotransferase | lacZ derived signal peptide | 2 | Fed-batch | 8 |
| PNGase F | ompA | 0.008 | Flask | 9 |
